# Skin-resident immune cells engulf axonal debris in adult epidermis

**DOI:** 10.1101/2022.06.15.496311

**Authors:** Eric Peterman, Elgene Quitevis, Emma C. Horton, Rune L. Aelmore, Ethan White, Alvaro Sagasti, Jeffrey P. Rasmussen

**Author notes:** **Corresponding author/lead contact:** JPR.

## Abstract

Somatosensory neurons extend enormous peripheral axons to the skin, where they detect diverse environmental stimuli. Somatosensory peripheral axons are easily damaged due to their small caliber and superficial location. Axonal damage results in Wallerian degeneration, creating vast quantities of cellular debris that phagocytes must remove to maintain organ homeostasis. The cellular mechanisms that ensure efficient clearance of axon debris from stratified adult skin are unknown. Here, we establish zebrafish scales as a tractable model to study axon degeneration in the adult epidermis. Using this system, we demonstrate that skin-resident immune cells known as Langerhans cells engulf the majority of axon debris. In contrast to immature skin, adult keratinocytes do not significantly contribute to debris removal, even in animals lacking Langerhans cells. Our study establishes a powerful new model for studying Wallerian degeneration and identifies a new function for Langerhans cells in maintenance of adult skin homeostasis following injury. These findings have important implications for pathologies that trigger somatosensory axon degeneration.

## Introduction

Skin is a dynamic organ that constantly replenishes its constituent cells during homeostasis and in response to injury. Skin provides protection to environmental insults by functioning both as a durable barrier and sensory organ. Dense networks of somatosensory axon endings arborize throughout the epidermis, the outermost layer of the skin, and detect a variety of stimuli, including pain, temperature, and itch (Handler and Ginty, 2021). Cutaneous injuries and wounds damage these fragile axons, which can trigger Wallerian degeneration (WD), a molecular program of axon degeneration (Coleman and Höke, 2020). WD leaves the neuronal soma intact, but the axon distal to the injury degenerates. WD creates a myriad of axonal debris fragments that must be removed to restore full functionality to the skin. This presents a particular challenge for the skin given the enormous size and complexity of cutaneous arbors (Wu et al., 2012).

Phagocytic cells engulf and degrade axon debris following WD, which allows for potential axon reinnervation and promotes tissue homeostasis by preventing inflammation. Typically, peripheral nerve damage triggers recruitment of professional phagocytes derived from the immune system, which infiltrate the distal nerve and phagocytose axon debris (Zigmond and Echevarria, 2019). Surprisingly, studies in the larval skin of *Drosophila melanogaster* and *Danio rerio* revealed that, rather than relying on macrophages or other immune cells, epidermal keratinocytes engulf and degrade nearly all cutaneous somatosensory neurite debris (Han et al., 2014; Rasmussen et al., 2015). A limitation of these models is that the larval epidermis of these animals contains only a mono-or bilayer of keratinocytes and lacks the diverse repertoire of immune cell types that appear later in vertebrate skin organogenesis (Botting and Haniffa, 2020). Thus, whether these models accurately reflect the cellular and molecular mechanisms involved in removal of axonal debris in mature, stratified skin remains unknown. Identification of the phagocytes involved in debris removal in the adult epidermis is relevant for understanding post-embryonic pathologies in which axon homeostasis is altered, such as diabetic and chemotherapy-induced peripheral neuropathy (Stucky and Mikesell, 2021).

Here, we develop an *ex vivo* model to assess somatosensory axon degeneration and subsequent phagocytosis in adult zebrafish scales. Our approach allows live-cell imaging of WD in the presence of all resident cell types found in the adult vertebrate epidermis, including stratified keratinocytes and diverse immune cell types. By imaging axon degeneration and the associated cellular responses, we identify the cells responsible for axon debris clearance following degeneration in adult epidermis. In contrast to larval animals, epidermal keratinocytes do not play a major role in debris engulfment. Rather, we find that Langerhans cells, a skin-resident immune cell type mainly studied for their antigen-presenting roles and not known to participate in skin repair (Kaplan, 2017), engulf the majority of cutaneous axon debris. Notably, keratinocytes do not engulf increased quantities of axonal debris in the absence of Langerhans cells. Altogether, our work establishes scale explants as a tractable system to study WD, allowing us to image this process in a stereotypical fashion with high spatiotemporal resolution. We specifically highlight the cell biology of injury responses in the adult skin and reveal that larval and adult skin use different mechanisms for debris removal.

## Results

### The adult zebrafish scale as a model for Wallerian degeneration

In order to follow the fate of degenerating cutaneous axons in adult skin, we sought a simple method to trigger WD of somatosensory axons. We previously demonstrated that the peripheral axons of dorsal root ganglion (DRG) somatosensory neurons densely innervate the epidermis above adult scales (Rasmussen et al., 2018). Scales are dermal appendages that cover the adult trunk in an overlapping pattern akin to tiles on a roof (Figure 1A). Physically removing (“plucking”) scales from adult fish removes both the bony scale and attached epidermis with resident cell types including keratinocytes and peripheral axons. Scale removal severs cutaneous axons from their somata (Figure 1A). To determine whether axons degenerate within explanted scales, we plucked and cultured scales from adults expressing a reporter for a subset of somatosensory neurons *(Tg(p2rx3a:lexA;LexAOP:mCherry),* hereafter referred to as *Tg(p2rx3a:mCherry);* (Palanca et al., 2013)). Using live-cell imaging to monitor axon degeneration in real-time (Figure 1A), we found that approximately half of the mCherry+ axons initiated degeneration between 165 and 240 minutes post-pluck, generating significant amounts of axon debris in the epidermis (Figures 1B,1E, and 1F; Supplemental Video 1).

**Figure 1.**
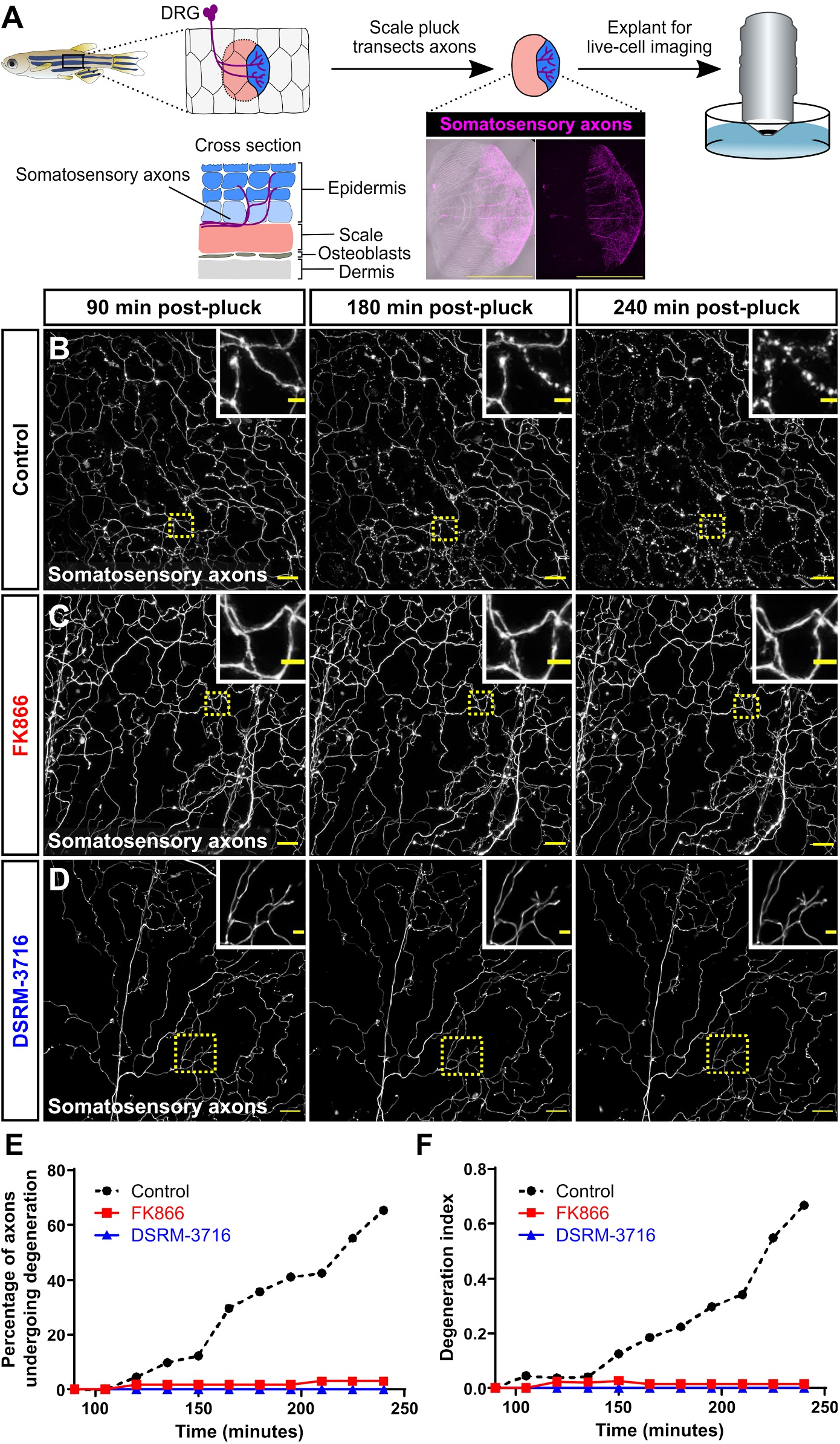
*ex vivo* scale explants as a model for Wallerian degeneration. (A) Schematic depicting the anatomy of the adult zebrafish scale epidermis and scale removal. The stratified epidermis (above the bony scale) is innervated by the peripheral axons of dorsal root ganglion somatosensory neurons. Confocal image shows an example of an entire scale explant expressing a somatosensory axon reporter (magenta; *Tg(p2rx3a:mCherry)).* (B-D) Confocal images of time-lapses of explanted scales from adults expressing a somatosensory axon reporter *(Tg(p2rx3a:mCherry)).* Scales were treated with DMSO as a vehicle control (B), FK866 (C), or DSRM-3716 (D). Dotted boxes denote the regions magnified in insets. Note the lack of axon degeneration in the FK866- and DSRM-3716- treated scales. See Supplemental Video 1. (E,F) Percentage of axons undergoing degeneration (E) and the axon degeneration index (F) in control, FK866- treated and DSRM-3716- treated scales. *n*=8 for control, *n*=6 for FK866, *n*=8 for DSRM-3716 (regions of interest). Scale bars: 1 mm (A), 20 μm (B-D), 5 μm (B-D, insets).

To determine if the axon degeneration we observed was specifically due to WD, we incubated scales in two different small molecule inhibitors of WD. First, we used FK866, an inhibitor of nicotinamide phosphoribosyltransferase (Nampt). FK866 has previously been shown to inhibit WD in culture and following axotomy of larval somatosensory neurons (Di Stefano et al., 2015). Second, we used DSRM-3716, a sterile-α and Toll/interleukin 1 receptor motif containing protein 1 (Sarm1) antagonist (Hughes et al., 2021). SARM1 is necessary for the normal progression of WD (Osterloh et al., 2012). We found that, compared to vehicle-treated controls, treating explanted scales with either of these inhibitors potently blocked axon degeneration, up to 4 hours post-pluck (Figures 1C–1F; Supplemental Video 1). Thus, explanted scales can serve as a simple model of WD in the adult epidermis. Furthermore, inhibition of Nampt or Sarm1 after axon severing is sufficient to inhibit WD.

### Creation of transgenic tools to monitor keratinocyte phagosomes in adult epidermis

What are the cell(s) responsible for engulfing axon debris following WD in adult skin? We previously demonstrated that together the two layers of larval keratinocytes (periderm and basal cells) engulf essentially all cutaneous axon debris (Rasmussen et al., 2015). Similarly, keratinocyte-like epidermal cells internalize and degrade neurite debris in larval *Drosophila* skin (Han et al., 2014). Thus, we began by examining keratinocyte contributions to debris removal in adult skin. The epidermis stratifies during post-larval growth, adding layers of suprabasal cells in between the periderm and basal cell layers (Guzman et al., 2013; Rangel-Huerta et al., 2021). In adult scales, somatosensory axons arborize throughout basal and suprabasal epidermal layers, often in direct contact with keratinocytes (Rasmussen et al., 2018).

To track the phagocytic ability of adult keratinocytes by live-cell imaging, we expressed EGFP-2xFYVE, which binds to a phospholipid enriched on phagosome membranes (Gillooly et al., 2000), in two overlapping keratinocyte populations. First, we used upstream regulatory sequences from the keratinocyte marker *krt4* to drive expression of EGFP-2xFYVE *(Tg(krt4:EGFP-2xFYVE))*. *Tg(krt4:EGFP-2xFYVE*) labeled spherical, phagosome-like compartments within keratinocytes predominantly in the periderm layer (Figures S1A and S1B). Second, we engineered a BAC with EGFP-2xFYVE in place of the start codon of ΔNp63 (*TgBAC(ΔNp63:EGFP-2xFYVE)*), which we found labeled similar structures of basal and suprabasal keratinocytes (Figures S1A and S1C). We further validated that these transgenes label keratinocyte phagosomes by demonstrating that the EGFP-2xFYVE+ structures co-localized with LysoTracker, a live-cell dye that labels acidic compartments (Figure S1D). Together these new tools allow for the unambiguous tracking of keratinocyte phagocytosis at larval and adult stages.

### Keratinocytes do not play a significant role in debris engulfment following axon degeneration

To assess keratinocyte involvement in axon debris clearance in adult epidermis, we live imaged scale explants with axons labeled with mCherry and *krt4+/ΔNp63+* keratinocyte phagosomes labeled with EGFP-2xFYVE. The relative pH-stability of mCherry (Cranfill et al., 2016) allowed us to track axon debris over extended periods of time. We found that, similar to larval keratinocytes, adult keratinocytes could internalize axon debris (Figure 2A, arrowheads). However, adult keratinocytes did not significantly contribute to debris removal, engulfing less than 10% of axonal debris after WD (Figure 2A, 2E; Supplemental Video 2). These results suggest that axon debris either remains largely unengulfed, or a different phagocytic cell type clears axon debris in adults.

**Figure 2.**
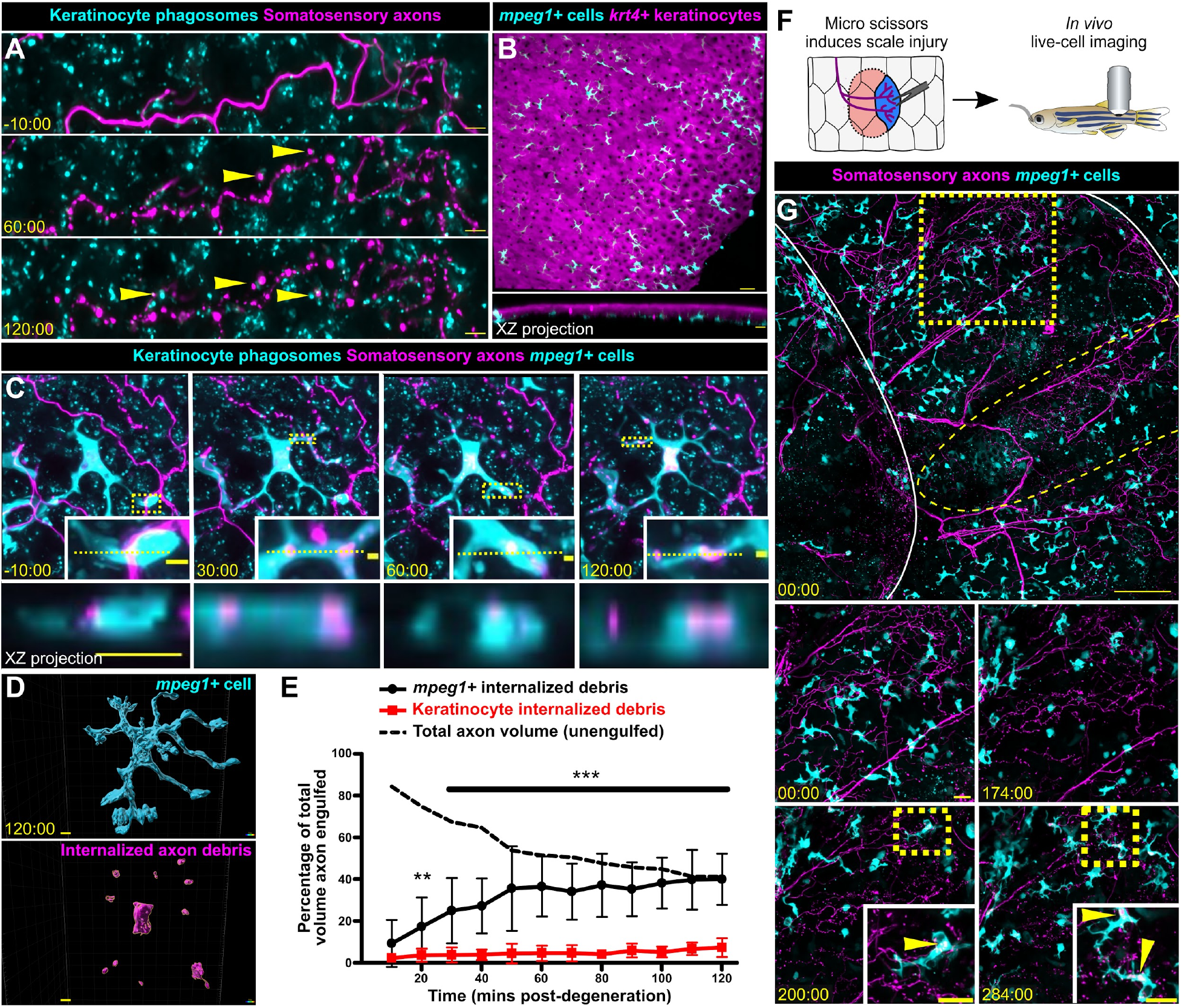
*mpeg1+* cells, not keratinocytes, engulf the majority of axon debris following axon degeneration. (A) Confocal images from a time-lapse of scale pluck-induced axon degeneration from an adult expressing reporters for keratinocyte phagosomes *(Tg(krt4:EGFP-2xFYVE); TgBAC(ΔNp63:EGFP-2xFYVE))* and somatosensory axons *(Tg(p2rx3a:mCherry)).* Arrowheads show examples of colocalization. Time denotes mm:ss relative to the onset of axon degeneration. See Supplemental Video 2. (B) Lateral confocal image (top) and reconstructed cross section (bottom) showing that *mpeg1+* cells (cyan) densely populate the scale epidermis, and reside beneath the *krt4+* layer (magenta). (C) Confocal images from a time-lapse of scale pluck-induced axon degeneration from an adult expressing reporters for keratinocyte phagosomes *(Tg(krt4:EGFP-2xFYVE);TgBAC(ΔNp63:EGFP-2xFYVE)),* somatosensory axons *(Tg(p2rx3a:mCherry)),* and *mpeg1+* cells *(Tg(mpeg1:NTR-EYFP))* before and during scale pluck-induced axon degeneration. Yellow dotted lines in insets denote the plane reconstructed in the XZ projections. Time denotes mm:ss relative to the onset of axon degeneration. See Supplemental Videos 3 and 4. (D) Surface view used in Imaris from panel C (120:00) for volume engulfed quantifications. (E) Quantification of total axon volume engulfed over time by keratinocytes and *mpeg1+* cells. Two-way ANOVA followed by Bonferroni tests determined significance between *mpeg1+ cells* and keratinocytes. ** = *p*<0.01, *** = *p*<0.001. *n*=10-16 cells/regions of interest. (F) Schematic for inducing scale injury *in vivo.* (G) Confocal images from a time-lapse of *in vivo* scale injury from an adult expressing reporters for somatosensory axons *(Tg(p2rx3a:mCherry)),* and *mpeg1+* cells *(Tg(mpeg1:NTR-EYFP).* Solid white lines outline scales from fish, yellow dotted oval denotes site of micro scissor injury, yellow dotted box denotes inset for bottom panels. Time denotes mm:ss relative to the time of injury. See Supplemental Videos 5 and 6. Scale bars: 5 μm (A,D), 30 μm (B), 10 μm (C), 2 μm (C, insets), 100 μm (G, upper panel), 20 μm (G, lower panels).

### mpeg1+ cells engulf large quantities of debris following axon degeneration

Since keratinocytes did not engulf appreciable amounts of axonal debris, we questioned if other cell types clear axonal debris in adult epidermis. Previous reports suggest a variety of immune cells populate the skin throughout organogenesis (Kasheta et al., 2017; Kuil et al., 2020; Lin et al., 2019; Lugo-Villarino et al., 2010; Wittamer et al., 2011); therefore, we reasoned that the adult epidermis would have a larger variety of immune cells present to possibly participate in debris engulfment. Using cell type-specific transgenic animals, we found that the adult scale epidermis contained *mpeg1+* Langerhans cells and metaphocytes, *lck+* lymphocytes, and *mpx+* neutrophils (Figure 2B, S2).

Due to the high density of *mpeg1+* cells and their roles in repair and phagocytosis in other contexts (Casano et al., 2016; Peri and Nüsslein-Volhard, 2008; Petrie et al., 2014; Rosenberg et al., 2012), we postulated they may participate in axon debris engulfment. To directly compare the relative contributions of *mpeg1+* cells and keratinocytes to debris removal, we created quadruple transgenic animals expressing reporters for *mpeg1+* cells, *krt4+/ΔNp63+* keratinocyte phagosomes, and axons. By live-cell imaging axon degeneration over a period of hours, we found that *mpeg1+* cells engulfed debris at a significantly higher efficiency in comparison to keratinocytes (Figures 2C–2E and Supplemental Video 3 and 4). Orthogonal cross sections revealed that debris was fully internalized within *mpeg1+* cells, which frequently relied on dynamic protrusions to internalize the debris (Figure 2C).

To further validate that our scale explant is an accurate model for observing cutaneous axon degeneration and engulfment, we asked whether *mpeg1+* cells engulfed axonal debris *in vivo.* To test this, we combined a scale injury model in adult fish with previous methods of intubation and imaging to observe Langerhans cells in living animals (Figure 2F). Axon degeneration was observed 2-4 hours after injury, consistent with the timing of Wallerian degeneration *ex vivo* (Figure 2G and Supplemental Videos 5 and 6). Following axon degeneration, we observed *mpeg1+* cells engulfing axonal debris (Figure 2G, insets and Supplemental Videos 5 and 6), providing evidence that *mpeg1+* cells clear cutaneous axonal debris *in vivo.* Together, these data strongly suggest that *mpeg1+* Langerhans cells and/or metaphocytes internalize axon debris following WD in adult epidermis.

### Langerhans cells constitute the mpeg1+ cell population that engulfs axon debris

As described above, *mpeg1* reporters label multiple cell types in the adult epidermis. This includes both Langerhans cells, a skin-resident dendritic cell known mainly for their antigen-presenting properties in mammals (Kaplan, 2017), and metaphocytes, a recently identified epidermal cell type in zebrafish (Alemany et al., 2018; Lin et al., 2019). A subset of dendritic cells isolated from adult zebrafish skin contain Birbeck granules (Lugo-Villarino et al., 2010), a defining characteristic of Langerhans cells (Birbeck et al., 1961; Valladeau et al., 2000), suggesting that zebrafish skin contains *bona fide* Langerhans cells. Interestingly, a previous report suggested that metaphocytes uptake soluble antigens, which they transfer to Langerhans cells via an apoptosis-phagocytosis mechanism (Lin et al., 2019).

To distinguish between possible contributions of Langerhans cells and metaphocytes to axonal debris engulfment, we took three parallel approaches. First, we used a previously described methodology based on morphological differences (Kuil et al., 2020). In this approach, Langerhans cells are distinguished by multiple, branched protrusions, while metaphocytes have fewer, less complex protrusions (Figure 3A). We quantified the volume of internalized debris within individual *mpeg1+ cells* at 60 minutes post-axon degeneration and found that debris engulfment positively correlated with protrusion number (Figure 3B), suggesting that the subset of *mpeg1+* cells we observed engulfing axonal debris were Langerhans cells. Second, we performed fluorescent *in situ* hybridization for *cd4-1,* a gene expressed by Langerhans cells but not metaphocytes (Lin et al., 2019). Scales were removed and fixed 3 hours post-removal, a time point at which axons are degenerating and Langerhans cells contain axon debris. *cd4-1* signal colocalized with *mpeg1+* cells that had internalized axon debris (Figure 3C). Third, we used a transgenic approach to differentially label Langerhans cells and metaphocytes during axon degeneration and debris engulfment. RNA-seq transcriptional profiling indicates that Langerhans cells, but not metaphocytes, express the gene *mfap4* (Kuil et al., 2020; Lin et al., 2019). To determine whether a previously generated *mfap4* transgene could be used to label Langerhans cells, we crossed *Tg(mfap4:tdTomato-CAAX);* (Walton et al., 2015) fish with *Tg(mpeg1:NTR-EYFP);Tg(p2rx3a:mCherry)* fish. In these triple transgenic animals, we expected Langerhans cells would be *mpeg1+/mfap4+,* whereas metaphocytes would be *mpeg1+/mfap4-.* Indeed, we found that *mfap4+* cells that were also *mpeg1+* had the highly branched morphology consistent with Langerhans cells, whereas metaphocytes remained only *mpeg1+* (Figure 3D). Removing scales and imaging for axonal debris engulfment revealed that *mpeg1+/mfap4+* cells specifically engulfed axonal debris (Figure 3E, F). Based on these results, we concluded that Langerhans cells are the primary adult cell type responsible for engulfing cutaneous axon debris.

**Figure 3.**
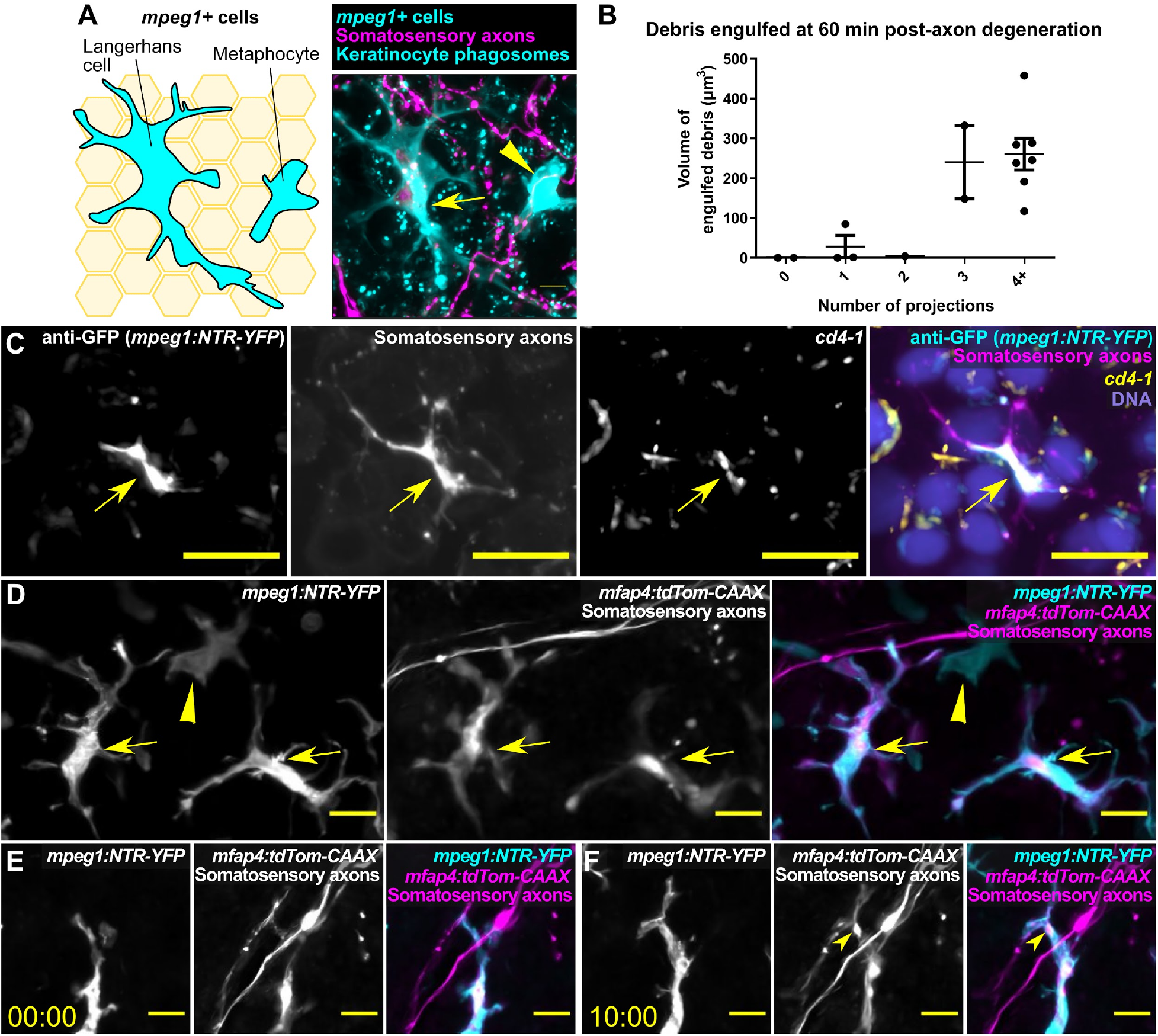
Langerhans cells represent the *mpeg1+* cell type that engulf cutaneous axon debris. (A) Schematic (left) and fluorescent image (right) comparing the morphology of a Langerhans cell (arrow) and a metaphocyte (arrowhead). (B) Quantification of axonal debris engulfed relative to number of protrusions in *mpeg1+* cells 60 minutes post scale pluck-induced axon degeneration. (C) Fluorescence *in situ* hybridization with an antisense probe against *cd4-1* following scale pluck-induced axon degeneration. Arrow indicates *cd4-1* expression in *mpeg1+* cell. (D) Fluorescence images of the scale epidermis in an adult expressing *Tg(mpeg1:NTR-EYFP), Tg(mfap4:tdTom-CAAX),* and a somatosensory axon reporter *(Tg(p2rx3a:mCherry)).* Arrows indicate *mpeg1+/mfap+* cells (Langerhans cells); arrowhead indicates an *mpeg1+* only cell (metaphocyte) (E, F) Still images of the scale epidermis in a *Tg(mpeg1:NTR-EYFP);Tg(mfap4:tdTom-CAAX);Tg(p2rx3a:mCherry)* adult before and during scale pluck-induced axon degeneration. Arrowhead indicates engulfed axonal debris. Time denotes mm:ss. Scale bars: 5 μm (A), 10 μm (C-F).

### Keratinocytes do not compensate for the loss of Langerhans cells

In direct contrast to larval skin, where keratinocytes are the primary phagocytes for axon debris (Han et al., 2014; Rasmussen et al., 2015), we found that adult keratinocytes largely do not contribute to debris clearance (Figure 2). Does the presence of highly phagocytic Langerhans cells in adult skin inhibit the ability of keratinocytes to engulf axon debris? To address this question, we used a previously described transgenic system *(Tg(mpeg1:NTR-EYFP);* (Petrie et al., 2014)) to conditionally ablate *mpeg1+* cells. *Tg(mpeg1:NTR-EYFP)* fish express the nitroreductase (NTR) enzyme fused to EYFP under the *mpeg1* promoter. Upon exposure to the prodrug metronidazole (MTZ), NTR converts MTZ into a toxic compound, thereby killing *mpeg1+* cells. We found that treating *Tg(mpeg1:NTR-EYFP)* adults with 7 mM MTZ for 3 days ablated most *mpeg1+* cells (Figure 4A, B). As controls, we treated siblings without *Tg(mpeg1:NTR-EYFP)* with MTZ. By removing scales and imaging axon degeneration following MTZ exposure, we observed only a small increase in axon engulfment by keratinocytes in the absence of *mpeg1+* cells (Figure 4C). As a parallel strategy, we next examined mutants that lack developmental colonization of the epidermis by Langerhans cells, a process that requires Colony stimulating factor-1 receptor (Csf1r) (Dai et al., 2002; Kuil et al., 2020). We first confirmed that animals with loss-of-function mutations in both of the zebrafish *csf1r* paralogs *(csf1ra^mh5/mh5^; csf1rb^mh108/mh108^;* (Caetano-Lopes et al., 2020)) lacked Langerhans cells (Figure 4D, E). Using *csf1ra^mh5/mh5^; csf1rb^mh108/mh108^* or double heterozygous animals, we repeated the engulfment assays and found that genetic ablation of Langerhans cells resulted in no change in the amount of debris engulfed by keratinocytes (Figure 4F). Thus, keratinocytes do not engulf more axon debris in the absence of Langerhans cells.

**Figure 4.**
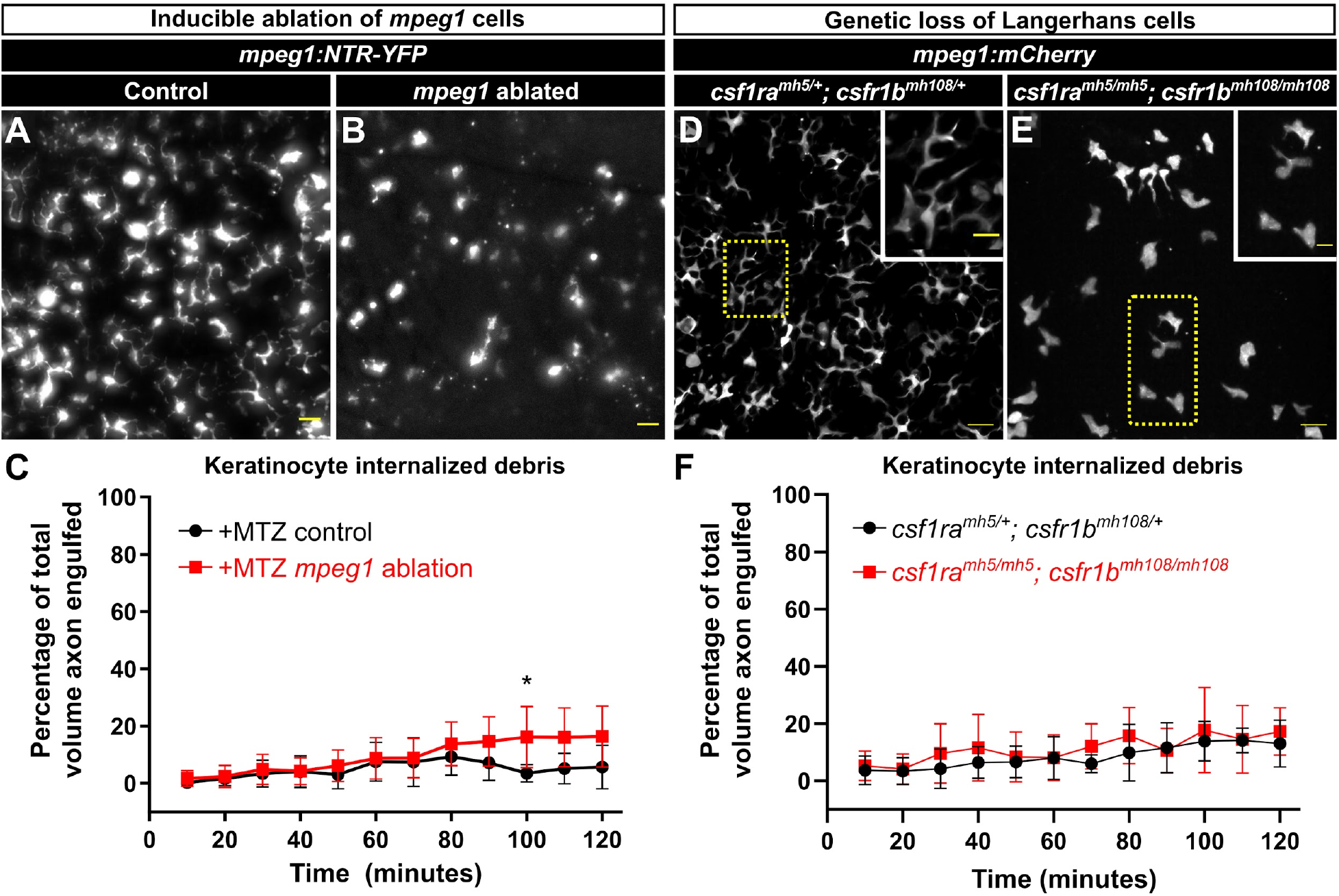
Keratinocytes do not compensate for the absence of Langerhans cells. (A,B) Representative widefield images of the scale epidermis in *Tg(mpeg1:NTR-EYFP)* adults after three days of mock treatment (A) or 7 mM MTZ treatment to ablate *mpeg1+* cells (B, 71% decrease in *mpeg1+* cells). (C) Quantification of debris engulfed by keratinocytes after three days of exposure to MTZ in animals with or without *Tg(mpeg1:NTR-EYFP)* (denoted as +MTZ *mpeg1* ablation or +MTZ control, respectively). Two-way ANOVA followed by Bonferroni tests determined significance between (+MTZ-NTR) and (+MTZ+NTR) conditions. * = *P* < 0.05. *n*=4- 10 regions of interest for +MTZ-NTR (from 3 fish, 6 scales) and *n*=15-16 regions of interest for +MTZ+NTR (from 7 fish, 13 scales) (D,E) Representative confocal images of *mpeg1+* cells in the scale epidermis from adults of the indicated genotypes. Note the lack of *mpeg1+* cells with the dendritic morphology of Langerhans cells in the *csf1ra^mh5/mh5^; csf1rb^mh108/mh108^* mutant epidermis (89% of *mpeg1+* cells have ≤ 1 protrusion). (F) Quantification of debris engulfed by keratinocytes in adults of the indicated genotypes. *n*=6-11 regions of interest for *csf1ra^mh5/+^; csf1rb^mh108/+^* (from 3 fish, 4 scales) and *n*=10 regions of interest for *csf1ra^mh5/mh5^; csf1rb^mh108/mh108^*(from 3 fish, 5 scales). Scale bars: 20 μm (A, B, D, E), 10 μm (D, E insets).

## Discussion

In this study, we combine a new scale explant technique with high-resolution confocal imaging to examine the mechanisms of axonal debris clearance following WD. In contrast to studies in larval models, we show that keratinocytes do not play a major role in axonal debris removal in adult epidermis. Instead, we demonstrate that Langerhans cells are the primary phagocytes responsible for the clearance of axonal debris following injury-induced WD. We observed that Langerhans cells do not prematurely attack axons, but rather engulf debris only after an axon has undergone WD. Combined, our data suggest differences in phagocytic properties in larval versus adult skin and reveal an unappreciated role for Langerhans cells in promoting skin homeostasis after injury.

We were surprised to find that adult keratinocytes have a greatly reduced phagocytic capability compared to their larval counterparts (Figure 2C). Several possible explanations exist for this apparent decline. First, the bilayered epidermis of larval zebrafish is much simpler in comparison to the stratified adult epidermis. A more constrained three-dimensional and/or adhesive environment could decrease the ability for keratinocytes to rearrange their plasma membranes in order to engulf debris. Meanwhile, Langerhans cells possess dynamic cellular protrusions (Nishibu et al., 2006), allowing them to navigate this complex environment to seek out and engulf axonal debris. A second possibility is that keratinocytes lose phagocytic competency through downregulation of the necessary machinery as organogenesis progresses. Controlled studies comparing larval and adult skin should be performed to fully investigate this scenario. Third, infiltration of immune cells, such as Langerhans cells, that are specialized for debris removal could trigger a change in keratinocyte gene expression. A study examining the transcriptomes of keratinocytes in the presence or absence of Langerhans cells revealed differential gene expression in keratinocytes (Su et al., 2020). However, our analysis of these results showed no upregulation of pro-engulfment genes in the absence of Langerhans cells, consistent with our observations that the presence of Langerhans cells does not inhibit keratinocyte phagocytosis (Figure 4).

Historically, Langerhans cells have been studied for their role as dendritic cells with antigen-presenting capabilities (Kaplan, 2017). After encountering foreign antigens in the skin, they drain to lymph nodes to interact with T cells, thereby regulating adaptive immune responses to cutaneous infections and allergens. Our work establishes a new role for Langerhans cells in maintaining skin homeostasis by engulfing cellular debris from degenerating axons, a function typically ascribed to tissue-resident macrophages. Future work will address the phagocytic pathways used by Langerhans cells during debris engulfment, including the machinery involved in recognizing, engulfing and degrading axonal debris. While not the focus of this study, an interesting future area of exploration will be to examine if Langerhans cells communicate with T cells to present antigen following WD, or whether Langerhans cells are solely responsible for the processing and degradation of axon debris without involvement of other cells.

**A** growing body of literature argues that Langerhans cells and microglia, tissue-resident macrophages of the central nervous system, share a number of characteristics. Langerhans cells and microglia arise from the same embryonic progenitor cells (Gomez Perdiguero et al., 2015) and both require interleukin-34 (Il-34) and Csf1r signaling for tissue infiltration (Kuil et al., 2020; Wang et al., 2012). Both cell types have dynamic cellular protrusions that survey the environment and transcriptomic studies indicate they have overlapping gene expression profiles (Mass et al., 2016). Intriguingly, our work shows that Langerhans cells are phagocytes of neuronal debris, similar to microglia in multiple vertebrate systems. Microglia have been well-characterized for their roles in synapse remodeling during development, where they can prune developing arbors according to neuronal activity (Bachiller et al., 2018). We hypothesize that Langerhans cells likely play similar roles in shaping cutaneous axon arbors during skin organogenesis.

In summary, we show that Langerhans cells clear axonal debris following WD in the epidermis. Of note, Langerhans cells ablation has been associated with a decrease in cutaneous axon density in mouse (Doss and Smith, 2014; Zhang et al., 2021), suggesting skin innervation or axon maintenance requires Langerhans cells. In addition, Langerhans cells may have roles in mediating or exacerbating peripheral neuropathies, conditions in which epidermal innervation is decreased, resulting in perturbations to skin sensation. Peripheral neuropathies such as chemotherapy-induced neuropathy and diabetic peripheral neuropathy result in altered numbers of Langerhans cells in the epidermis (Siau et al., 2006; Stojadinovic et al., 2013). The relationship between Langerhans cells and these neuropathies is currently poorly understood, but is of potential clinical significance. Recent work suggests that Langerhans cells mediate pain by directly signaling to sensory neurons (Raymondi Silva et al., 2022). Our findings could have relevance for identifying mechanisms that mediate the initiation or progression of peripheral neuropathies and future studies in zebrafish could provide a unique perspective on the role of Langerhans cells in these human diseases.

## Acknowledgements

We thank the LSB Aquatics staff for animal care and the labs of Matthew Harris, Shuo Lin, Cressida Madigan, Randall Moon, and David Tobin for sharing reagents. The authors are grateful to all members of the Rasmussen lab for discussion, technical assistance, and continuous support.

## Materials and methods

### Key resources table

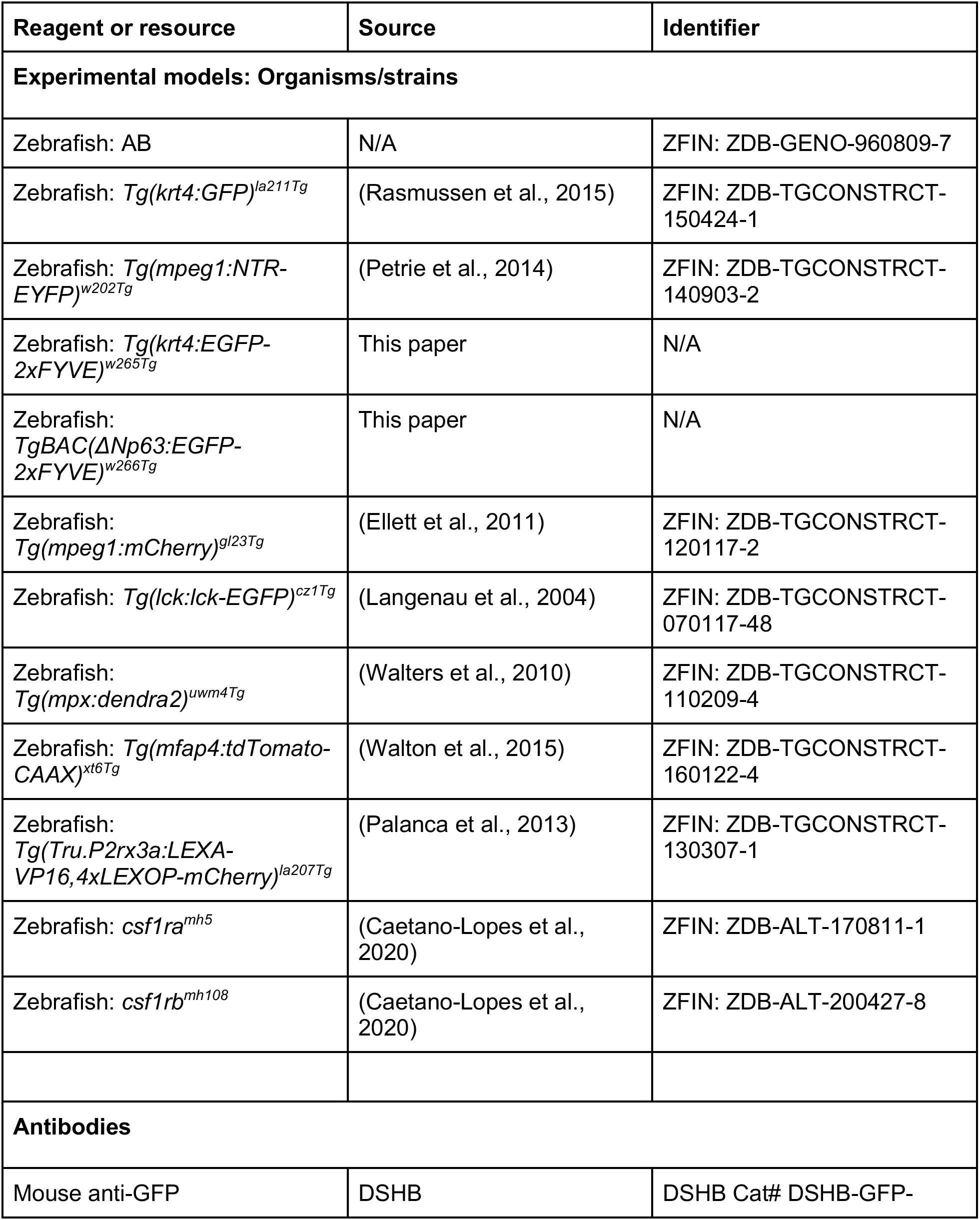
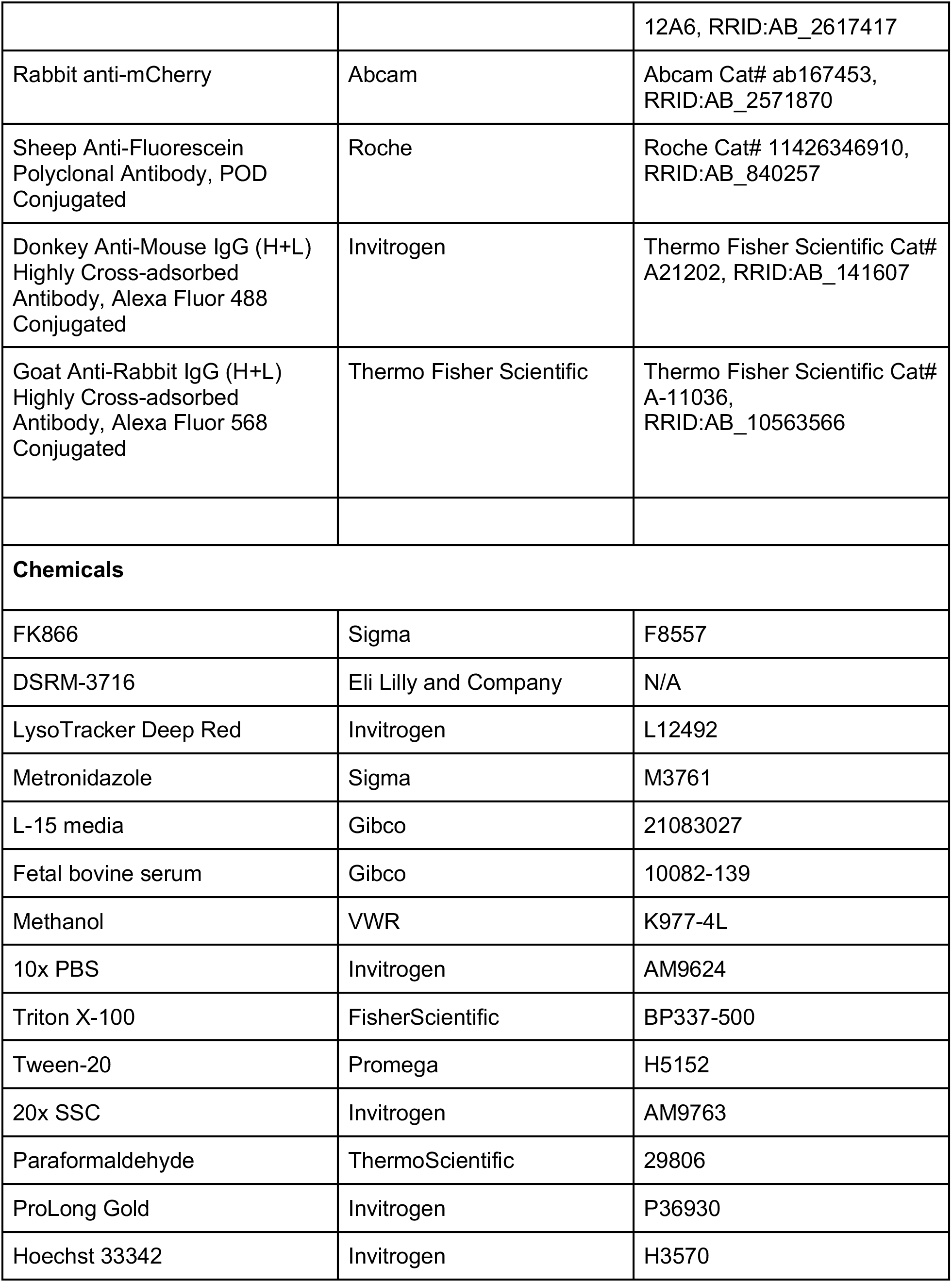
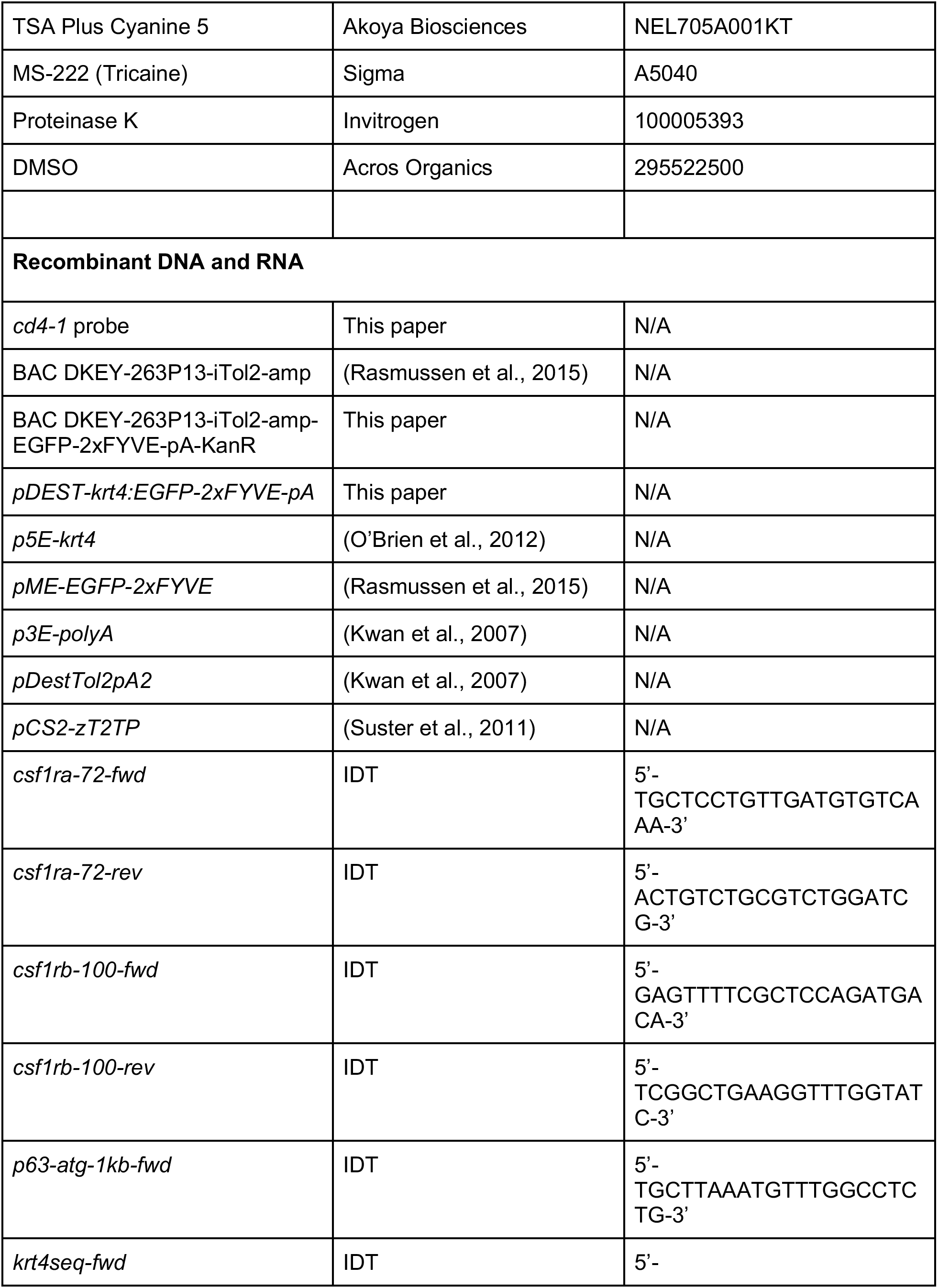
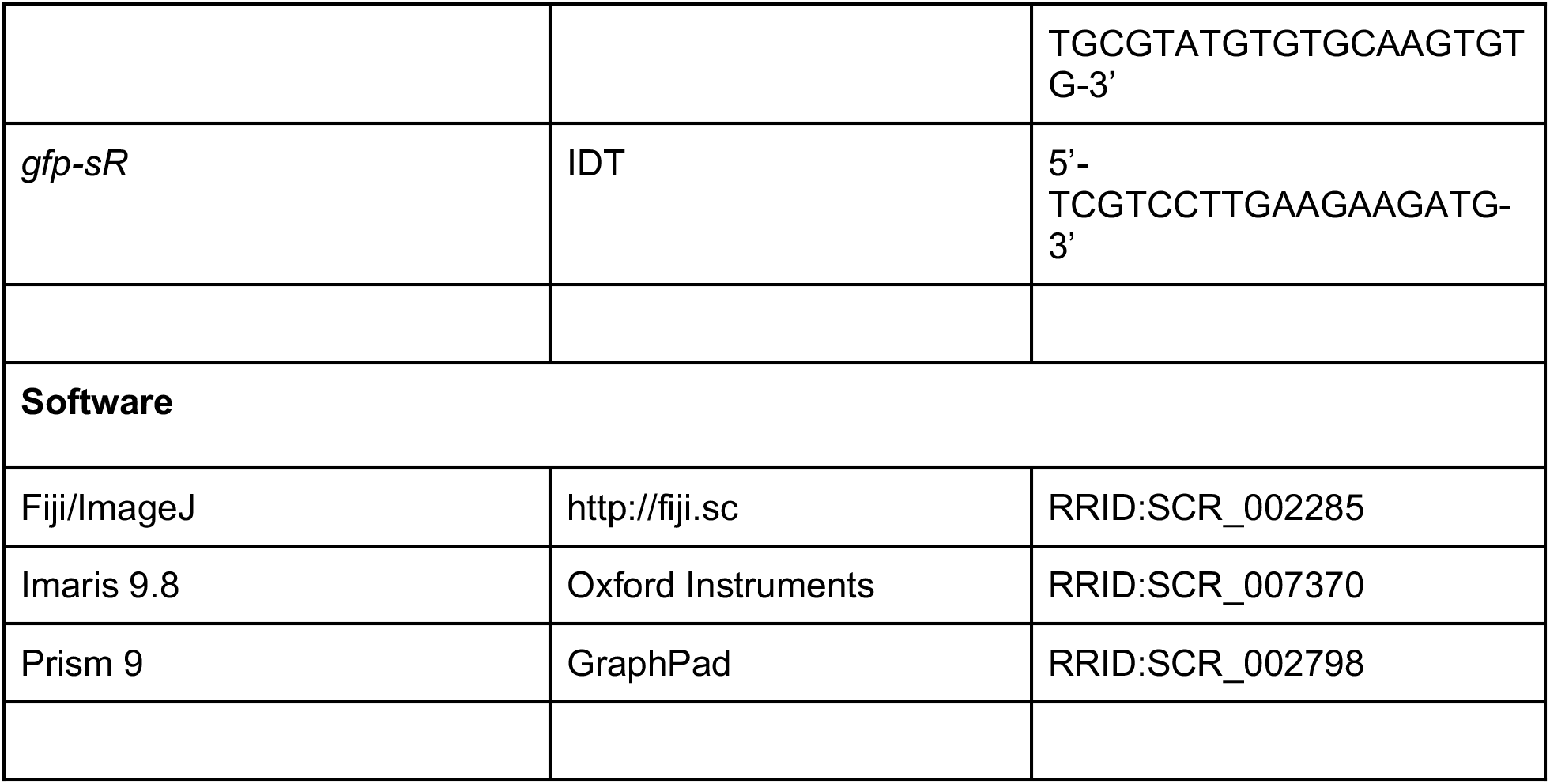

### Resource availability

#### Lead contact

Further information and request for resources and reagents should be directed and will be fulfilled by the Lead Contact; Jeff Rasmussen.

#### Materials availability

All unique/stable reagents generated in this study are available from the lead contact without restriction.

### Experimental model and subject details

#### Zebrafish and husbandry

Zebrafish were housed at 26-27°C on a 14/10 h light cycle. The strains used are referenced in the Key resources table. Animals of either sex were used in this study. All zebrafish experiments were approved by the Institutional Animal Care and Use Committee at the University of Washington (Protocol: #4439-01).

### Method details

#### Genotyping

Adult fish were genotyped to ensure that both *Tg(krt4:EGFP-2xFYVE)* and *Tg(ΔNp63:EGFP-2xFYVE)* were present. The primer pairs used were *krt4seq-fwd/gfp-sR* and *p63-atg-1kb-fwd/gfp-sR* to detect *Tg(krt4:EGFP-2xFYVE)* and *Tg(ΔNpβ3:EGFP-2xFYVE),* respectively. The *csf1ra^mh5^* and *csf1rb^mh108^* alleles were followed with high-resolution melt analysis using the primer pairs *csf1ra-72-fwd/csf1ra-72-rev* and *csf1rb-100-fwd/csf1rb-100-rev,* respectively.

#### Transgene construction

The *ΔNp63:EGFP-2xFYVE* bacterial artificial chromosome (BAC) was created by modifying the previously generated BAC DKEY-263P13-iTol2-amp (Rasmussen et al., 2015). The predicted *ΔNp63* start codon was replaced by a *EGFP-2xFYVE-pA-KanR* cassette using a previously described protocol (Suster et al., 2011). The *pDEST-krt4:EGFP-2xFYVE-pA* plasmid was assembled using Gateway recombination of *p5E-krt4* (O’Brien et al., 2012), *pME-EGFP-2xFYVE* (Rasmussen et al., 2015), *p3E-polyA,* and *pDestTol2pA2* (Kwan et al., 2007). *Tg(krt4:EGFP-2xFYVE)^w265Tg^* and *TgBAC(ΔNp63:EGFP-2xFYVE)^w266Tg^* were created by injecting *tol2* mRNA, which was transcribed from pCS2-zT2TP (Suster et al., 2011), and either plasmid or BAC DNA into one-cell stage embryos and screening adults for germline transmission.

#### Scale removal

For scale removal, adult fish were anesthetized in system water containing 200 μg/ml Tricaine and forceps were used to remove individual scales. Following scale removal, animals were recovered in system water.

#### Microscopy and live-imaging

An upright Nikon Ni-E A1R MP+ confocal microscope was used for all experiments. A 25x water dipping objective (NA 1.1) was routinely used. Unless otherwise stated, scales were removed and placed onto dry 6 mm plastic dishes, epidermis side up, and allowed to adhere for 1 minute before adding L-15 media pre-warmed to room temperature. Scales were incubated at 26°C for 90 min followed by imaging, which was performed at room temperature (23°C).

#### In vivo scale injury and imaging

For *in vivo* injury and imaging, adult fish were anesthetized in system water containing 200 μg/ml Tricaine. Sterile micro scissors were used to cut through a scale on the lateral trunk. Fish were subsequently immobilized and mounted in a custom chamber using 1% agarose dissolved in system water. Fish were intubated with aerated system water containing 120 μg/ml Tricaine. The chamber was placed under a 16x water dipping objective (NA 0.8), and the site of injury was located. Imaging was performed using an environmental imaging enclosure set to 28.5°C. After the imaging session, fish were recovered with aerated system water. Fish were placed back into system water following complete recovery (tail fin movement and rapid gill movement).

#### Chemical treatments and live-cell staining

For FK866 and DSRM-3716 treatments, scales were removed and immediately placed in L-15 containing 10 μM FK866 or 10 μM DSRM-3716. Scales were incubated for 90 min at 26°C before imaging commenced.

For metronidazole treatments, fish were placed in system water containing 7 mM metronidazole for 3 days. Fish were housed at a density of 1 fish per liter. Metronidazole solution was replaced daily.

For LysoTracker staining, scales were removed and placed in a 1.5 ml tube containing 2 nM LysoTracker Deep Red. Scales were incubated in the dark for 30 min at 26°C, washed quickly in L-15, and placed onto a 6 mm dish, epidermis side up. Scales were incubated in L-15 at 26°C in the dark for a further 60 min before imaging commenced.

#### Image analysis

To quantify percentage axon degeneration in Figure 1, random ROIs were selected and tracked every 15 min. The total axon number was used to calculate the percentage of axons undergoing degeneration. Degeneration index was calculated as previously described (Sasaki et al., 2009). Briefly, images were thresholded in ImageJ, and total axon intensity and degenerated axon intensity were calculated. The degeneration index represents the fraction of degenerated axon intensity over the total axon intensity.

To quantify debris engulfment, the Imaris Surfaces function was used. Surfaces were created for Langerhans cells, keratinocyte EGFP-2xFYVE+ phagosomes, and axons. Additional Surfaces were created using the same intensity threshold as the axon Surface, then filtered to only include material inside the “Langerhans cell” Surface or “FYVE” Surface. Images were manually inspected and corrected to ensure no erroneous material was counted as inside a Surface. Volumes for total axon volume and engulfed volume were recorded every 10 min and percentage debris engulfed was calculated.

To differentiate metaphocytes from Langerhans cells in Figure 3, the number of cellular protrusions on *mpeg1+* cells were counted at 60 min post-axon degeneration. A protrusion was defined as a process that extended ?5 microns from the cell body.

#### Fluorescent in situ hybridization (FISH)

A 350 bp gBlock fragment for the 3’ coding region of *cd4-1* was synthesized and inserted into pCS2+. Antisense RNA was transcribed *in vitro* using SP6 and fluorescein-dUTP. The FISH protocol for adult zebrafish scales was previously described (Lin et al., 2019). Briefly, scales from *Tg(mpeg1:NTR-EYFP);Tg(p2rx3a:mCherry)* adults were plucked and incubated in L-15 at 28°C for 3 h to ensure axon degeneration. Scales were fixed in 4% PFA overnight at 4°C then washed three times with 1xPBS + 0.1% Tween-20 (PBST). Scales were dehydrated in sequential washes of 75%PBST:25% methanol (MeOH), 50%PBST:50%MeOH, 25%PBST:75%MeOH, then placed in 100% MeOH at-20°C overnight. Scales were rehydrated in sequential washes of 25%PBST:75%MeOH, 50%PBST:50%MeOH, 75%PBST:25% MeOH, then washed 3x in PBST. Scales were treated with 0.1mg/ml proteinase K for 5 min, then re-fixed in 4% PFA for 20 min. Scales were washed once in PBST, washed once in 50%PBST:50% hybridization buffer, then incubated in hybridization buffer for 2 h at 65°C. Scales were incubated in hybridization buffer with *cd4-1* probe (~1 ng/μl) overnight at 65°C. Scales were sequentially washed at 65°C in 75% hybridization buffer:25% 2xSSC +0.1%Tween20 (SSCT), 50% hybridization buffer:50% 2xSSCT, 25% hybridization buffer:75% 2xSSCT, followed by 3 washes at room temperature in 2xSSCT, followed by 3 washes in 0.2xSSCT. Scales were then washed 3x in PBST, then blocked for 2 h in PBST+5%FBS. Scales were incubated in blocking buffer with anti-fluorescein POD fragments (1:2000) overnight at 4°C. Scales were washed 6x in PBST, followed by staining with TSA Plus Cyanine 5 (1:50 dilution) for 10 min.

Following FISH, scales were incubated in PBST+10%FBS for 2 h at room temperature. Scales were stained with anti-GFP (1:10) and anti-mCherry (1:50) antibodies in PBST+10%FBS overnight at 4°C. Scales were washed in PBST, then incubated in secondary antibodies (1:200) for 2 h at room temperature. Scales were washed in PBST, stained with Hoechst (3.24 nM) for 20 min at room temperature, washed in PBST, mounted under coverslips in ProLong Gold, and subsequently imaged.

#### Statistical analysis

GraphPad Prism was used to generate graphs and perform statistical analyses. Tests used and number of animals, scales or cells/ROIs are described in each figure legend.

## Supplemental Materials

**Supplemental Figure 1.**
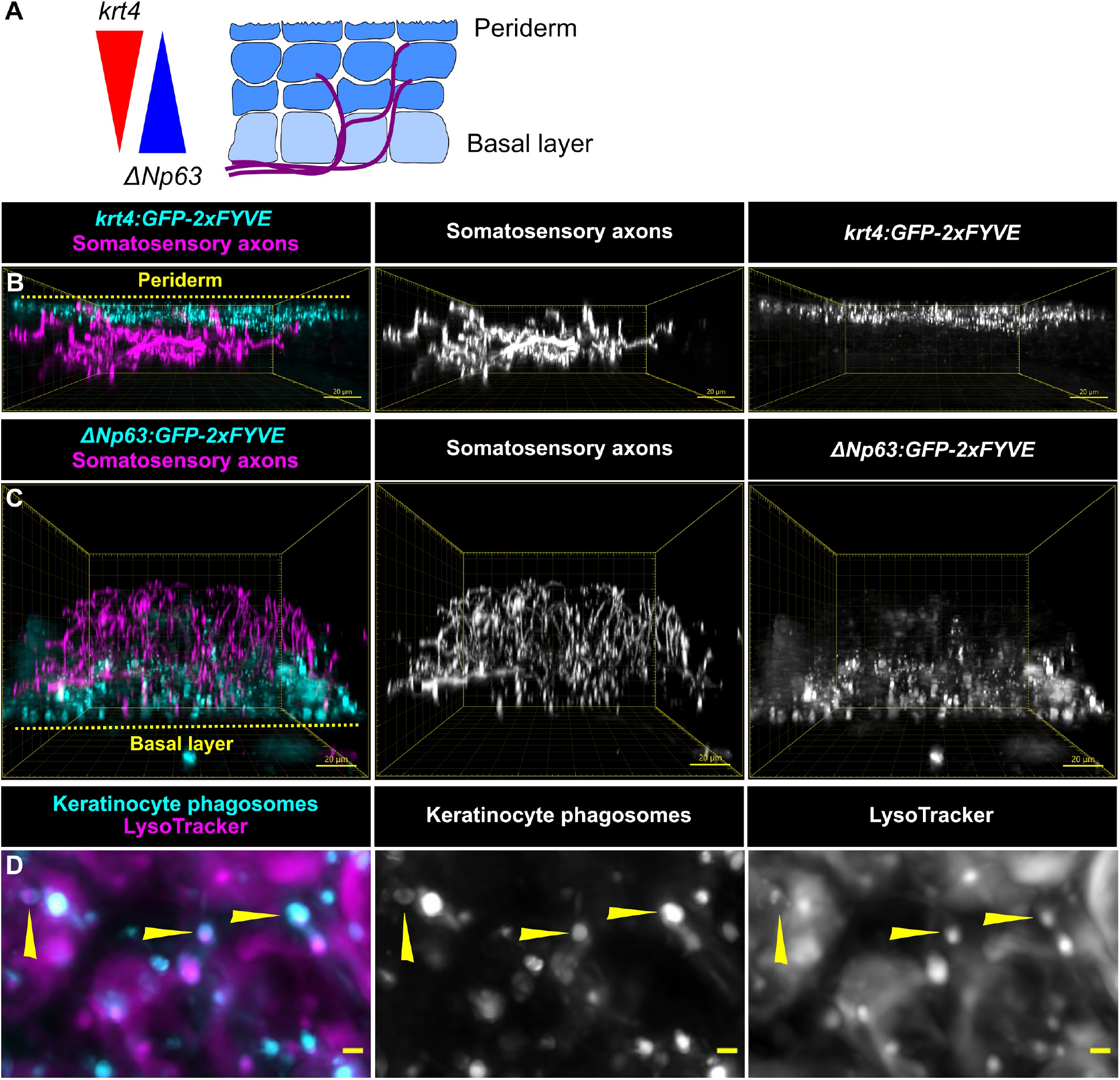
Characterization of epidermal keratinocyte transgenes. (A) Schematic depicting gradients of transgene expression patterns in adult epidermis. (B, C) Reconstructed cross-sections showing expression of *Tg(krt4:EGFP-2xFYVE)* and *TgBAC(ΔNp63:EGFP-2xFYVE)* in adult epidermis. (D) LysoTracker co-localization with EGFP-2xFYVE+ structures (arrowheads). Scale bars: 10 μm (B,C), 1 μm (D).

**Supplemental Figure 2.**
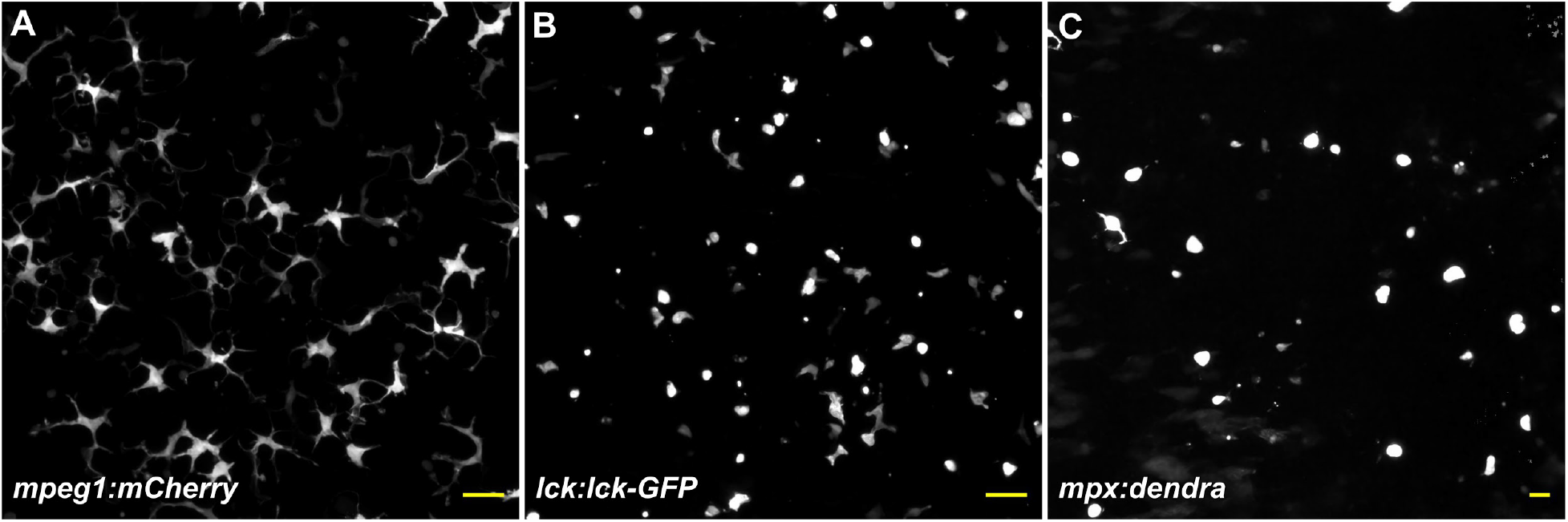
Identification of different immune cell populations present in adult epidermis. (A-C) Lateral confocal views of the adult scale epidermis expressing the indicated transgenes. Scale bars: 20 μm.

**Video 1**. Time-lapse microscopy of *Tg(p2rx3a:mCherry)+* axons in control conditions or treated with FK866 or DSRM-3716.

**Video 2**. Time-lapse microscopy of *Tg(krt4:EGFP-2xFYVE);TgBAC(ΔNp63:EGFP-2xFYVE)+* phagosomes engulfing a small quantity of *Tg(p2rx3a:mCherry)+* axon debris. Time 0 denotes onset of axon degeneration.

**Video 3**. Time-lapse microscopy of *Tg(p2rx3a:mCherry)+* debris being internalized by *Tg(mpeg1:NTR-EYFP)* cells. Time 0 denotes onset of axon degeneration.

**Video 4.** Imaris volume-blended time-lapse microscopy of *Tg(p2rx3a:mCherry)+* debris being internalized by *Tg(mpeg1:NTR-EYFP)* cells.

**Video 5**. *In vivo* time-lapse microscopy of a scale injury in *Tg(p2rx3a:mCherry;Tg(mpeg1:NTR-EYFP)* animals.

**Video 6**. Cropped and magnified view from Video 5 showing *Tg(p2rx3a:mCherry)+* debris being internalized by *Tg(mpeg1:NTR-EYFP)* cells.

## References

Alemany, A., Florescu, M., Baron, C. S., Peterson-Maduro, J. and van Oudenaarden, A. (2018). Whole-organism clone tracing using single-cell sequencing. Nature 556, 108–112.

Bachiller, S., Jiménez-Ferrer, I., Paulus, A., Yang, Y., Swanberg, M., Deierborg, T. and Boza-Serrano, A. (2018). Microglia in Neurological Diseases: A Road Map to Brain-Disease Dependent-Inflammatory Response. Front. Cell. Neurosci. 12,.

Birbeck, M. S., Breathnach, A. S. and Everall, J. D. (1961). An electron microscope study of basal melanocytes and high-level clear cells (Langerhans cells) in vitiligo. J. Invest. Dermatol. 37, 51–64.

Botting, R. A. and Haniffa, M. (2020). The developing immune network in human prenatal skin. Immunology 160, 149–156.

Caetano-Lopes, J., Henke, K., Urso, K., Duryea, J., Charles, J. F., Warman, M. L. and Harris, M. P. (2020). Unique and non-redundant function of csf1r paralogues in regulation and evolution of post-embryonic development of the zebrafish. Dev. Camb. Engl. 147, dev181834.

Casano, A. M., Albert, M. and Peri, F. (2016). Developmental Apoptosis Mediates Entry and Positioning of Microglia in the Zebrafish Brain. Cell Rep. 16, 897–906.

Coleman, M. P. and Höke, A. (2020). Programmed axon degeneration: from mouse to mechanism to medicine. Nat. Rev. Neurosci. 21, 183–196.

Cranfill, P. J., Sell, B. R., Baird, M. A., Allen, J. R., Lavagnino, Z., de Gruiter, H. M., Kremers, G.-J., Davidson, M. W., Ustione, A. and Piston, D. W. (2016). Quantitative assessment of fluorescent proteins. Nat. Methods 13, 557–562.

Dai, X.-M., Ryan, G. R., Hapel, A. J., Dominguez, M. G., Russell, R. G., Kapp, S., Sylvestre, V. and Stanley, E. R. (2002). Targeted disruption of the mouse colony-stimulating factor 1 receptor gene results in osteopetrosis, mononuclear phagocyte deficiency, increased primitive progenitor cell frequencies, and reproductive defects. Blood 99, 111–120.

Di Stefano, M., Nascimento-Ferreira, I., Orsomando, G., Mori, V., Gilley, J., Brown, R., Janeckova, L., Vargas, M. E., Worrell, L. A., Loreto, A., et al. (2015). A rise in NAD precursor nicotinamide mononucleotide (NMN) after injury promotes axon degeneration. Cell Death Differ. 22, 731–742.

Doss, A. L. N. and Smith, P. G. (2014). Langerhans cells regulate cutaneous innervation density and mechanical sensitivity in mouse footpad. Neurosci. Lett. 578, 55–60.

Ellett, F., Pase, L., Hayman, J. W., Andrianopoulos, A. and Lieschke, G. J. (2011). mpeg1 promoter transgenes direct macrophage-lineage expression in zebrafish. Blood 117, e49–56.

Gillooly, D. J., Morrow, I. C., Lindsay, M., Gould, R., Bryant, N. J., Gaullier, J. M., Parton, R. G. and Stenmark, H. (2000). Localization of phosphatidylinositol 3-phosphate in yeast and mammalian cells. eMbO J. 19, 4577–4588.

Gomez Perdiguero, E., Klapproth, K., Schulz, C., Busch, K., Azzoni, E., Crozet, L., Garner, H., Trouillet, C., de Bruijn, M. F., Geissmann, F., et al. (2015). Tissue-resident macrophages originate from yolk-sac-derived erythro-myeloid progenitors. Nature 518, 547–551.

Guzman, A., Ramos-Balderas, J. L., Carrillo-Rosas, S. and Maldonado, E. (2013). A stem cell proliferation burst forms new layers of P63 expressing suprabasal cells during zebrafish postembryonic epidermal development. Biol Open 2, 1179–1186.

Han, C., Song, Y., Xiao, H., Wang, D., Franc, N. C., Jan, L. Y. and Jan, Y.-N. (2014). Epidermal cells are the primary phagocytes in the fragmentation and clearance of degenerating dendrites in Drosophila. Neuron 81, 544–560.

Handler, A. and Ginty, D. D. (2021). The mechanosensory neurons of touch and their mechanisms of activation. Nat. Rev. Neurosci. 22, 521–537.

Hughes, R. O., Bosanac, T., Mao, X., Engber, T. M., DiAntonio, A., Milbrandt, J., Devraj, R. and Krauss, R. (2021). Small Molecule SARM1 Inhibitors Recapitulate the SARM1-/-Phenotype and Allow Recovery of a Metastable Pool of Axons Fated to Degenerate. Cell Rep. 34, 108588.

Kaplan, D. H. (2017). Ontogeny and function of murine epidermal Langerhans cells. Nat. Immunol. 18, 1068–1075.

Kasheta, M., Painter, C. A., Moore, F. E., Lobbardi, R., Bryll, A., Freiman, E., Stachura, D., Rogers, A. B., Houvras, Y., Langenau, D. M., et al. (2017). Identification and characterization of T reg-like cells in zebrafish. J. Exp. Med. 214, 3519–3530.

Kuil, L. E., Oosterhof, N., Ferrero, G., Mikulášová, T., Hason, M., Dekker, J., Rovira, M., van der Linde, H. C., van Strien, P. M., de Pater, E., et al. (2020). Zebrafish macrophage developmental arrest underlies depletion of microglia and reveals Csf1r-independent metaphocytes. eLife 9, e53403.

Kwan, K. M., Fujimoto, E., Grabher, C., Mangum, B. D., Hardy, M. E., Campbell, D. S., Parant, J. M., Yost, H. J., Kanki, J. P. and Chien, C.-B. (2007). The Tol2kit: a multisite gateway-based construction kit for Tol2 transposon transgenesis constructs. Dev. Dyn. Off. Publ. Am. Assoc. Anat. 236, 3088–3099.

Langenau, D. M., Ferrando, A. A., Traver, D., Kutok, J. L., Hezel, J.-P. D., Kanki, J. P., Zon, L. I., Look, A. T. and Trede, N. S. (2004). In vivo tracking of T cell development, ablation, and engraftment in transgenic zebrafish. Proc. Natl. Acad. Sci. U. S. A. 101, 7369–7374.

Lin, X., Zhou, Q., Zhao, C., Lin, G., Xu, J. and Wen, Z. (2019). An Ectoderm-Derived Myeloid-like Cell Population Functions as Antigen Transporters for Langerhans Cells in Zebrafish Epidermis. Dev. Cell 49, 605–617.e5.

Lugo-Villarino, G., Balla, K. M., Stachura, D. L., Bañuelos, K., Werneck, M. B. F. and Traver, D. (2010). Identification of dendritic antigen-presenting cells in the zebrafish. Proc. Natl. Acad. Sci. U. S. A. 107, 15850–15855.

Mass, E., Ballesteros, I., Farlik, M., Halbritter, F., Günther, P., Crozet, L., Jacome-Galarza, C. E., Händler, K., Klughammer, J., Kobayashi, Y., et al. (2016). Specification of tissue-resident macrophages during organogenesis. Science 353, aaf4238.

Nishibu, A., Ward, B. R., Jester, J. V., Ploegh, H. L., Boes, M. and Takashima, A. (2006). Behavioral Responses of Epidermal Langerhans Cells In Situ to Local Pathological Stimuli. J. Invest. Dermatol. 126, 787–796.

O’Brien, G. S., Rieger, S., Wang, F., Smolen, G. A., Gonzalez, R. E., Buchanan, J. and Sagasti, A. (2012). Coordinate development of skin cells and cutaneous sensory axons in zebrafish. J. Comp. Neurol. 520, 816–831.

Osterloh, J. M., Yang, J., Rooney, T. M., Fox, A. N., Adalbert, R., Powell, E. H., Sheehan, A. E., Avery, M. A., Hackett, R., Logan, M. A., et al. (2012). dSarm/Sarm1 is required for activation of an injury-induced axon death pathway. Science 337, 481–484.

Palanca, A. M. S., Lee, S.-L., Yee, L. E., Joe-Wong, C., Trinh, L. A., Hiroyasu, E., Husain, M., Fraser, S. E., Pellegrini, M. and Sagasti, A. (2013). New transgenic reporters identify somatosensory neuron subtypes in larval zebrafish. Dev. Neurobiol. 73, 152–167.

Peri, F. and Nüsslein-Volhard, C. (2008). Live Imaging of Neuronal Degradation by Microglia Reveals a Role for v0-ATPase a1 in Phagosomal Fusion In Vivo. Cell 133, 916–927.

Petrie, T. A., Strand, N. S., Yang, C.-T., Tsung-Yang, C., Rabinowitz, J. S. and Moon, R. T. (2014). Macrophages modulate adult zebrafish tail fin regeneration. Dev. Camb. Engl. 141, 2581–91.

Rangel-Huerta, E., Guzman, A. and Maldonado, E. (2021). The dynamics of epidermal stratification during post-larval development in zebrafish. Dev. Dyn. Off. Publ. Am. Assoc. Anat. 250, 175–190.

Rasmussen, J. P., Sack, G. S., Martin, S. M. and Sagasti, A. (2015). Vertebrate epidermal cells are broad-specificity phagocytes that clear sensory axon debris. J. Neurosci. Off. J. Soc. Neurosci. 35, 559–70.

Rasmussen, J. P., Vo, N.-T. and Sagasti, A. (2018). Fish scales dictate the pattern of adult skin innervation and vascularization. Dev. Cell 46, 344–359.e4.

Raymondi Silva, J., Iftinca, M., Fernandes Gomes, F. I., Segal, J. P., Smith, O. M. A., Bannerman, C. A., Silva Mendes, A., Defaye, M., Robinson, M. E. C., Gilron, I., et al. (2022). Skin-resident dendritic cells mediate postoperative pain via CCR4 on sensory neurons. Proc. Natl. Acad. Sci. 119, e2118238119.

Rosenberg, A. F., Wolman, M. A., Franzini-Armstrong, C. and Granato, M. (2012). In Vivo Nerve-Macrophage Interactions Following Peripheral Nerve Injury. J. Neurosci. 32, 3898–3909.

Sasaki, Y., Vohra, B. P. S., Lund, F. E. and Milbrandt, J. (2009). Nicotinamide Mononucleotide Adenylyl Transferase-Mediated Axonal Protection Requires Enzymatic Activity But Not Increased Levels of Neuronal Nicotinamide Adenine Dinucleotide. J. Neurosci. 29, 5525–5535.

Siau, C., Xiao, W. and Bennett, G. J. (2006). Paclitaxel-and vincristine-evoked painful peripheral neuropathies: Loss of epidermal innervation and activation of Langerhans cells. Exp. Neurol. 201, 507–514.

Stojadinovic, O., Yin, N., Lehmann, J., Pastar, I., Kirsner, R. S. and Tomic-Canic, M. (2013). Increased number of Langerhans cells in the epidermis of diabetic foot ulcers correlates with healing outcome. Immunol. Res. 57, 222–228.

Stucky, C. L. and Mikesell, A. R. (2021). Cutaneous pain in disorders affecting peripheral nerves. Neurosci. Lett. 765, 136233.

Su, Q., Bouteau, A., Cardenas, J., Uthra, B., Wang, Y., Smitherman, C., Gu, J. and Igyártó, B. Z. (2020). Brief communication: Long-term absence of Langerhans cells alters the gene expression profile of keratinocytes and dendritic epidermal T cells. PLOS ONE 15, e0223397.

Suster, M. L., Abe, G., Schouw, A. and Kawakami, K. (2011). Transposon-mediated BAC transgenesis in zebrafish. Nat. Protoc. 6, 1998–2021.

Valladeau, J., Ravel, O., Dezutter-Dambuyant, C., Moore, K., Kleijmeer, M., Liu, Y., Duvert-Frances, V., Vincent, C., Schmitt, D., Davoust, J., et al. (2000). Langerin, a novel C-type lectin specific to Langerhans cells, is an endocytic receptor that induces the formation of Birbeck granules. Immunity 12, 71–81.

Walters, K. B., Green, J. M., Surfus, J. C., Yoo, S. K. and Huttenlocher, A. (2010). Live imaging of neutrophil motility in a zebrafish model of WHIM syndrome. Blood 116, 2803–2811.

Walton, E. M., Cronan, M. R., Beerman, R. W. and Tobin, D. M. (2015). The Macrophage-Specific Promoter mfap4 Allows Live, Long-Term Analysis of Macrophage Behavior during Mycobacterial Infection in Zebrafish. PloS One 10, e0138949.

Wang, Y., Szretter, K. J., Vermi, W., Gilfillan, S., Rossini, C., Cella, M., Barrow, A. D., Diamond, M. S. and Colonna, M. (2012). IL-34 is a tissue-restricted ligand of CSF1R required for the development of Langerhans cells and microglia. Nat. Immunol. 13, 753–760.

Wittamer, V., Bertrand, J. Y., Gutschow, P. W. and Traver, D. (2011). Characterization of the mononuclear phagocyte system in zebrafish. Blood 117, 7126–7135.

Wu, H., Williams, J. and Nathans, J. (2012). Morphologic diversity of cutaneous sensory afferents revealed by genetically directed sparse labeling. eLife 1, e00181.

Zhang, S., Edwards, T. N., Chaudhri, V. K., Wu, J., Cohen, J. A., Hirai, T., Rittenhouse, N., Schmitz, E. G., Zhou, P. Y., McNeil, B. D., et al. (2021). Nonpeptidergic neurons suppress mast cells via glutamate to maintain skin homeostasis. Cell 184, 2151–2166.e16.

Zigmond, R. E. and Echevarria, F. D. (2019). Macrophage biology in the peripheral nervous system after injury. Prog. Neurobiol. 173, 102–121.

